# Vasohibin-1 mediated tubulin detyrosination selectively regulates secondary sprouting and lymphangiogenesis in the zebrafish trunk

**DOI:** 10.1101/2020.04.23.053256

**Authors:** Bastos de Oliveira Marta, Meier Katja, Coxam Baptiste, Geudens Ilse, Jung Simone, Szymborska Anna, Gerhardt Holger

## Abstract

Previous studies have shown that Vasohibin-1 (Vash-1) is stimulated by VEGFs in endothelial cells and that its overexpression interferes with angiogenesis *in vivo*. Recently, Vasohibin-1 was found to mediate tubulin detyrosination, a post-translational modification that is implicated in many cell functions, such as cell division. Here we used the zebrafish embryo to investigate the cellular and subcellular mechanisms of Vash-1 on endothelial microtubules during formation of the trunk vasculature. We show that microtubules within venous-derived secondary sprouts are strongly and selectively detyrosinated in comparison with other endothelial cells, and that this difference is lost upon *vash-1* knockdown. Vasohibin-1 depletion in zebrafish specifically affected secondary sprouting from the posterior cardinal vein, increasing both the number of sprouts and endothelial cell divisions. We show that altering secondary sprout numbers and structure upon *vash-1* depletion leads to a failure in the development and specification of lymphatic vessels of the zebrafish trunk.

**SUMMARY:** Vasohibin-1 mediated detyrosination of endothelial microtubules is selectively required for adequate behaviour of venous secondary sprouting and subsequent formation of functional lymphatics in the zebrafish trunk.

## INTRODUCTION

Blood vessel formation and patterning is essential for tissue growth and homeostasis in vertebrate development and physiology. Endothelial cells arising from the lateral plate mesoderm in early embryonic development initially coalesce at the midline to form the first arterial and venous structures. Subsequent sprouting angiogenesis and remodelling expands and shapes the vascular tree throughout the developing embryo (Hogan & Schulte-Merker, 2017; Potente et al., 2011). Complex morphogenic and cell differentiation processes orchestrate formation of arteries, veins, and capillaries, as well as of the lymphatic vascular system.

The trunk vasculature in zebrafish embryos has served both angiogenesis and lymphangiogenesis research as a powerful model to observe, manipulate and mechanistically understand relevant cellular processes and their molecular control (Gore et al., 2016; Hogan & Schulte-Merker, 2017). Following the initial assembly and lumenisation of the dorsal aorta (DA) and posterior cardinal vein (PCV), the intersegmental vessels (ISV) form by sprouting. This first wave of sprouting emerges from the DA at 22 hours post fertilization (hpf), with bilateral sprouts along the somite boundaries to form the first vascular loops (Isogai et al., 2003; Kohli et al., 2013). A second wave of sprouting emerges from the PCV at 34hpf (Koltowska et al., 2015; Nicenboim et al., 2015; Yaniv et al., 2006), remodelling some ISVs into veins (Geudens et al., 2010; Yaniv et al., 2006) and forming parachordal lymphangioblasts (PLs) at the midline. The PLs subsequently migrate and branch to shape the mature lymphatic system of the zebrafish trunk including the thoracic duct (TD) and the dorsal longitudinal lymphatic vessel (DLLV) (Küchler et al., 2006; Yaniv et al., 2006). Although primary and secondary sprouts appear morphologically similar, with tip and stalk cells and acto-myosin cellular protrusions, they are driven by distinct growth factors and downstream signalling cascades. In particular, arterial sprouting involves VEGF-A, VEGFR2, PLCγ, and on the other hand venous/lymphatic sprouting is controlled by VEGF-C, VEGFR3, CCBE (Hogan & Schulte-Merker, 2017).

Recent reports have highlighted the importance of actin-based in endothelial cell (EC) dynamics during morphogenesis (Gebala et al., 2016; Lamalice et al., 2007; Phng et al., 2015). Microtubules however, although critically involved in cell division (Belmont et al., 1990), polarity (Siegrist & Doe, 2007) and vesicle transport mechanisms (Vale, 2003), have received little attention in vascular biology research. This is despite their relevance for tumour angiogenesis and vessel maintenance, as microtubule targeting drugs used for cancer therapy are anti-angiogenic and/or stimulate the collapse of tumour vasculature (Hayot et al., 2002; Kruczynski et al., 2006; Schwartz, 2009; Shi et al., 2016; Tozer et al., 2002). The mechanisms behind this phenomenon are not understood. Microtubules are unbranched cytoskeleton polymers made of α- and β-tubulin heterodimers that assemble in a directional manner to shape a polarised hollow tube. α-tubulin carries a tyrosine residue that can be removed post-translationally by the carboxylpeptidase Vasohibin-1 (Vash-1) (Aillaud et al., 2017; Nieuwenhuis et al., 2017), in a reaction referred to as tubulin detyrosination. Tubulin detyrosination is crucial for neuronal development (Erck et al., 2005), cell division (Barisic et al., 2015; Liao et al., 2019) and influences vesicle transport (Dunn et al., 2008). Interestingly, *VASH-1* was previously identified as a VEGFA-inducible gene *in vitro* (Abe & Sato, 2001) and is expressed in the developing endothelium *in vivo* (Hosaka et al., 2009; Shibuya et al., 2006; Watanabe et al., 2004). Viral overexpression as well as VASH-1 protein administration experiments *in vitro* and *in vivo* show that VASH-1 inhibits angiogenesis and reduces vessel diameter (Hosaka et al., 2009; Kern et al., 2009). Conversely, *VASH-1* knockout mice exhibited more vascularised tumours (Hosaka et al., 2009) and increased angiogenesis in the ear skin injury model (Kimura et al., 2009). Although these studies demonstrated that VASH-1 is necessary for controlling angiogenesis in a tumour and injury onset *in vivo*, they predated the discovery of VASH-1 as a microtubule modifying enzyme. Thus, the role of microtubule detyrosination and VASH-1 in particular in the context of *in vivo* angiogenesis remains unknown.

Here, we use the zebrafish trunk vasculature to characterise microtubule detyrosination and the effects of *vash-1* knockdown on secondary sprouting and lymphatic development. Using high-resolution live imaging, combined with immunofluorescence and gene expression analysis, we find that endothelial tubulin is predominantly detyrosinated in secondary sprouts. *vash-1* knockdown causes morphogenic defects in secondary but not primary sprouts, impeding proper lymphatic development. Mechanistically, *vash-1* loss-of-function leads to increased EC division within secondary sprouts and supernumerous secondary sprouts that fail to adopt lymphatic lineage identity. We propose that Vash-1 controlled endothelial tubulin detyrosination supports differential endothelial specification during cell division and sprout elongation to achieve venous and lymphatic vessel formation in the zebrafish trunk.

## MATERIAL AND METHODS

### Zebrafish maintenance and transgenic lines

Zebrafish (*Danio rerio*) were raised and staged as previously described (Kimmel et al., 1995). For growing and breeding of transgenic lines, we complied with regulations of the animal ethics committees at MDC Berlin (Aleström et al., 2019). The following transgenic lines were used: *Tg[fli1a:EGFP]*_*y1*_ (Lawson & Weinstein, 2002) labels all ECs, *Tg[fli1a:nEGFP]y7* labels all EC nuclei (Roman et al., 2002), *Tg[kdr-l:ras-Cherry]*_*s916*_ (Hogan, Bos, et al., 2009) labels EC membrane, *Tg[fli1ep:EGFP-DCX]* (Phng et al., 2015) labels endothelial doublecortin, a microtubule associated protein.

### Immunofluorescence staining and imaging

Embryos were treated with PTU in later stages than 24hpf to prevent pigment formation. Embryos were fixed with 4% Paraformaldehyde (PFA) for 10h at 4°C, and kept for a maximum of 4 days in PBTX at 4°C. Embryos were permeabilised by dehydrating them in a crescent methanol concentrations solutions - 25%, 50% and 75% -, incubated for 1h at −20°C, before being rehydrated in descending methanol concentration solutions - 75%, 50% and 25% -, 3 times washed with PBTX and post-fixed with 4% PFA for 10 minutes (min). The embryos were incubated for 2 hours in blocking buffer (10% horse heat inactivated serum, 1% DMSO, 100mM maleic acid in 0,1% PBTX) at room temperature. Embryos were incubated at 4°C overnight with primary antibodies (indicated below) diluted in blocking buffer. The embryos were then washed with a solution containing 1% DMSO and 100mM maleic acid in 0,1% PBTX at room temperature and blocked again for 30 min before being incubated with secondary antibodies diluted in blocking buffer overnight. Embryos were washed again in 1% DMSO and100mM maleic acid in 0,1% PBTX, fixed with PFA 4% for 30 min at room temperature, washed again and stored at 4 C in PBST until being mounted in glycerol and stored at 4°C.

Primary antibodies used were anti-detyrosinated tubulin (1:600, rabbit, home-made (Liao et al., 2019), anti-GFP FITC conjugate (1:200, mouse, Abcam, ref. ab6662). Secondary antibody used was Invitrogen Alexa 568 against rabbit (1:200 donkey, ref. A10037). All antibodies were diluted in the blocking buffer, described in the previous paragraph.

### Live imaging and analysis

Embryos were dechorionated and anaesthetised with 0,014% (1x) tricaine methanesulfonate (MS-222, Sigma-Aldrich). Embryos were then mounted in 0,8% low melting point agarose (Life Technologies) containing 0,014% tricaine and immersed in E3 buffer containing 0,014 % tricaine and 0,003% PTU when indicated. Live embryos were imaged with an upright 3i Spinning Disk Confocal using Zeiss Plan-Apochromat 20x and 40x /1.0 NA water dipping objectives. Image processing was performed using Fiji software (Schindelin et al., 2012). XY drifts were corrected using the MultiStackReg plugin (B. Busse, NICHD).

### Secondary sprout and 3-way connection parameters

The number of secondary sprouts was quantified in 15 minute time-lapse movies from 24 to 48 hpf, in *Tg[kdrl-l:ras-Cherry,fli1a:nEGFP]* embryos. The number of nuclei was assessed in each secondary sprouts just prior to connection to the pre-existing ISV. Cell divisions in secondary sprouts were quantified by adding nuclear (nEGFP) divisions from the time of emergence from the PCV, until the resolution of the 3-way connection with pre-existing ISVs. The duration of the 3-way connection was quantified in the same embryos from the moment it connects to the primary ISV and lumenises until the 3-way connection is resolved.

### Lymphatics: PLs and TD quantifications

PLs and TD were quantified by examining Z-stacks of 52 hpf and 4 days post fertilization (dpf) *Tg[fli1a:EGFPy1]* embryos and counting presence/absence in each somite. The number of PLs associated with venous ISVs, identified by a connection to the venous ISV, were identified at 52 hpf in *vash-1* morphants and control siblings.

### Immunofluorescence quantifications

Substacks containing single ISVs were cropped manually from images of whole *Tg[fli1ep:EGFP-DCX]* embryos stained with dTyr antibody. To estimate the background, which varied locally due to autofluorescence of the embryos, each image was filtered with a large Gaussian filter (radius = 50 pixels) and subtracted from the original image. A maximum intensity projection of the DCX channel was used to manually segment endothelial microtubules, with particular care to exclude the neurons. The resulting mask was applied to background corrected, sum intensity projections of DCX and dTyr channels. Mean intensity was extracted and plotted, to quantify the normalised and direct differences of tubulin detyrosination intensity in different experimental groups. The analysis was performed in Fiji.

### Statistical analysis

Zebrafish embryos were selected on the following pre-established criteria: normal morphology, beating heart, and flowing blood. The experiments were not randomised and the investigators were not blinded during experiment and analysis. All morphant embryos analysed in this project exhibited no apparent visual phenotype.

In order to decide if a parametric or non-parametric test should be used to compare different experimental groups, we assessed whether the population was normally distributed, by Anderson-Darling test. If so, t-test and regular ANOVA were performed, if not, Mann-Whitney and Kruskall-Wallis tests were performed.

### Cell isolation for gene expression analysis

*Tg[fli1a:nEGFP]*_*y7*_ were outcrossed and their embryos collected. Negative embryos were also collected under the stereoscope, to serve as GFP negative control siblings during cell sorting. Embryos were treated with 0,003% PTU when older than 24hpf. The embryos were dechorionated by adding pronase (1mg/ml) for 15 min, subsequently washed in E3 buffer, and anaesthetised with 0,16 mg/mL tricaine methanesulfonate (Sigma). 250 embryos were transferred to a 1.5ml tube and the yolk sac was removed by 2 rounds of centrifugation (or until the solution was clear) with Calcium ringer solution at 2000rpm at 4°C for 5 min. The embryos were then resuspended in 1ml of Protease solution (0,07mg/ml of Liberase in DPBS-Thermofisher) with 0,4U/ml DNAse I (Invitrogen). The embryos were dissociated at 28°C for 10min on a rotator (or until the embryos appeared visually digested to a single cell solution), pipetting up and down with a 200 mL tip every 4-5 min. The reaction was stopped by placing the cell solution on ice and adding CaCl2 to a final concentration of 1-2mM with 0,5% FBS in DPBS. The cells were centrifuged at 2000rpm for 5 min at 4°C, the supernatant was discarded, and the pellet resuspended in 1ml resuspension solution (DPBS with 2mM EDTA, 0,4U/ml DNAseI and 0,5%FBS) to avoid cells clumping together. The cells were passed through a 40μM strainer into to a 50ml falcon tub, following a pre-wash of the strainer with 500μl of the resuspension solution. Another 500μL of the resuspension solution was added to the filter to remove any cells left. The cells were centrifuged at 2000rpm for 5 min at 4°C and resuspended in 500μl of DPBS+EDTA solution. The cell solution was transferred to a 5ml polypropylene (Falcon) dedicated test tube and kept on ice until sorting with a FACSAria™ III Sorter (BD). Cells were sorted and filtered against triplets and droplets. Negative control cells from negative nEGFP siblings were used to define the GFP+ collection window, to filter against auto-fluorescence. nEGFP positive and negative cells from green *Tg[fli1a:nEGFP]*_*y7*_ embryos were collected in different 1.5ml eppendorf tubes with 350μl Trizol.

### RNA extraction

Cell lysis suspension was vortexed for 15 seconds (s), spun down, and incubated at room temperature for 5 min. 0,5μl glycoblue, a nucleic acid coprecipitant was added to the lysed solution to aid visualising the precipitated RNA. 50μl chloroform was added to each tube, which were shaken vigorously for 15 seconds, and incubated at room temperature for 5 min. Tubes were centrifuged at 12.000RPM for 15 min at 4°C. RNA is present in the transparent supernatant. The supernatant was carefully pipetted to a LoBind eppendorf tube. 125μl was added to the aqueous phase and the tube was shaken vigorously for 15s. The solution was incubated at −20°C overnight. The solution was then centrifuged at 12.000RPM for 10 min for 4°C and a small blue RNA pellet is visible at this point. The supernatant was then removed, leaving the RNA pellet that was washed with 500μl of 75% ethanol, shaken and centrifuged at 7.500RPM for 5 min at 4°C. The supernatant was discarded and the pellet air dried until no liquid was left. The pellet was resuspended in 13μl of DEPC water and the concentration and purity was measured in a Nanodrop 2000 (ThermoScientific) and the reverse transcription proceeded for samples presenting with a an RNA concentration above 20ng/μl and a 260/280 purity higher than 1.8. The kit Superscript IV First-Strand cDNA synthesis reaction (ThermoFisher) was used for the reverse transcription reaction, using random hexamer primers. The reaction was incubated for 1 hour. The final reaction was diluted 1:10 and kept at −20°C.

### qPCR

The primers were designed using Primer blast (http://www.ncbi.nlm.nih.gov/tools/primer-blast/) to have Tm of 60°C and to span exon-exon junctions to avoid amplification of genomic DNA. Each reaction was set in a 20μL volume, mix containing 10μl of SYBR-green qPCR mix (Roboklon), 0,8μl cDNA (1:50 dilution), 0,45μl of carboxyrhodamine (ROX) and 0,25μl uracil-N-glycosylase (UNG) from Roboklon and primers to a final concentration of 300uM, three technical replicates of each sample were added. Amplification was performed in triplicates in 384 well plates (Applied Biosystems) with the following thermal cycling conditions: initial UDG treatment 50°C for 10 min, followed by 40 cycles of 15 s at 95°C and 60 s at 60°C. Control reactions included a no template control (NTC). qPCR was performed in Quantstudio6Flex machine (Life-technologies). Amplification curves were analysed to confirm the presence of a single PCR product, and outliers excluded when one of the Ct values was very different from the other two. qPCR data was analysed by using the Pfaff method (Pfaffl, 2001) that integrates primer amplification efficiency, determined by LinReg software (Ruijter et al., 2009). Samples were normalised against 4 endothelial housekeeping genes-*rpl13*, *rps29*, *hprt1* and *ef2a*. The geometric average of the housekeeping genes was used for normalisation of gene expression (Coxam et al., 2014).

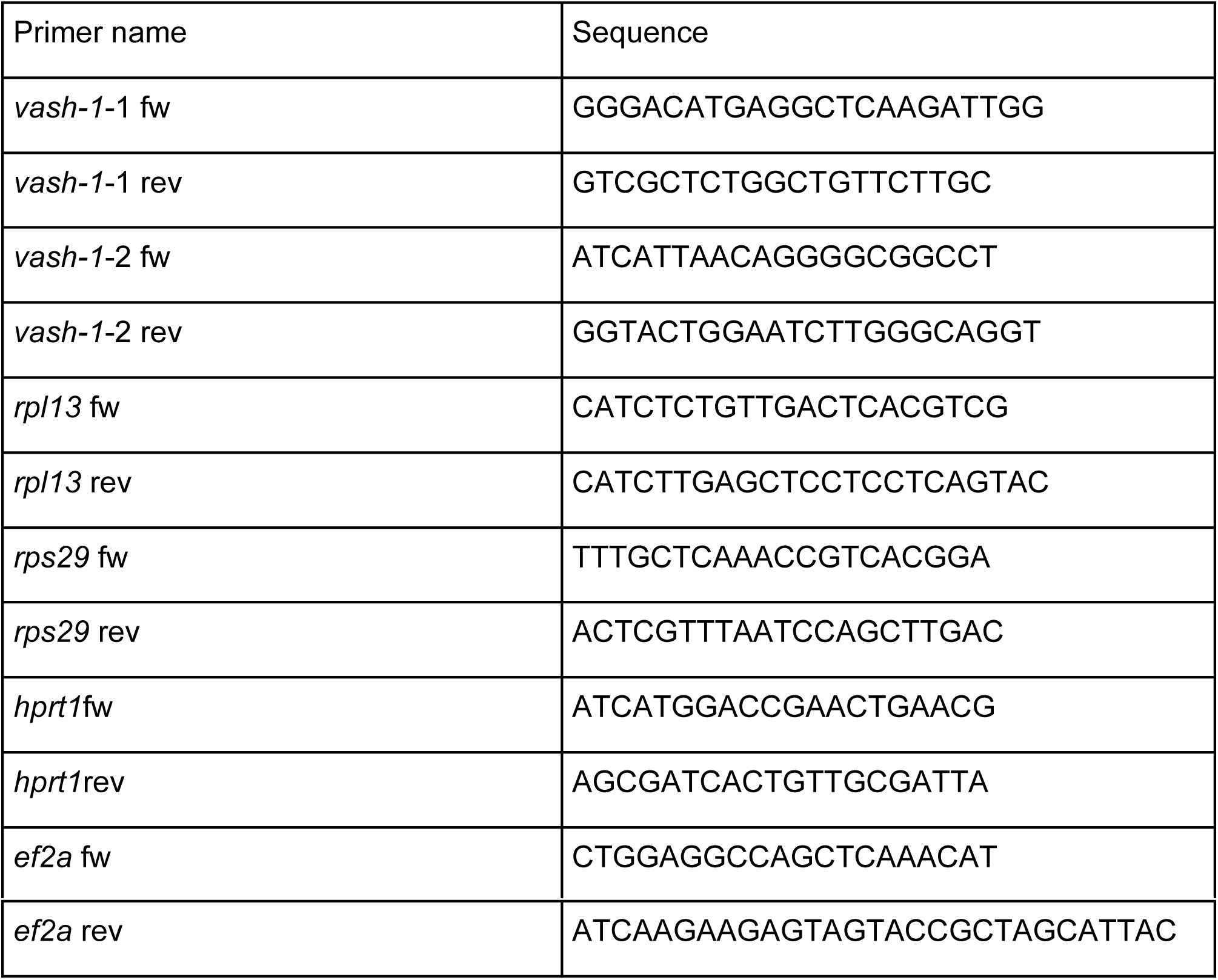

### Protein extraction for protein analysis

35 embryos were dechorionated by adding pronase (1mg/ml) and anaesthetised with 0,16 mg/mL tricaine methanesulfonate (Sigma). Yolk sacs were removed with 2 rounds of centrifugation in Calcium ringer solution at 2000rpm, 4°C, for 5 min and removed all liquid until only a dry pellet of embryos is left in the tube. Pellets were fast frozen with liquid nitrogen and kept at −80°C for at least 20min. Protein extraction procedure included adding 40 μl of lysis buffer (1M Tris-HCl,0,5M EDTA, 10% Brij 96, 10% NP-and 0,4 μl of protease inhibitor cocktail (ThermoFisher), homogenise it with a pestle (Starlab), centrifugation at 4°C for 15 min. The concentration of the proteins extracts were measured with the BCA protein assay kit (ThermoFisher).

### Western Blot

50μg of the lysates were mixed with reducing sample buffer (1:4) and heated for 5 min at 95°C to denaturate proteins. The protein were electrophoresised for 1h at 150V and subsequently transferred onto MtOH activated polyvinylidene fluoride membranes (GE Healthcare). Equal loading was assessed using Ponceau red solution. Membranes were blocked with Blocking solution (5% nonfat dry milk in TBS-T) for 1,5h at room temperature and then incubated with primary antibody against Vash-1, overnight at 4°C (1:500, Proteintech, ref. 12730-1-AP). After incubation with primary antibody, the membranes were washed 4 times in TBS-T and then incubated with secondary antibodies for 1h (1:4,000; GE Healthcare) and washed 3 times with TBS-T. Immunodetection was performed using a chemiluminescence kit (SuperSignal West Dura; Pierce), and bands were developed using the Las-4000 imaging system. After initial immunodetection, membranes were stripped of antibodies by using the Stripping kit (ThermoFisher) at 56°C for 40min and re-probed with anti– beta-actin antibody for 1h (1:1000, Sigma, ref. A5441).

### Morpholino experiments

Morpholino against *plcγ* was used as previously described (Hogan, Bos, et al., 2009). Dosage curves were used to find the appropriate dosage for injection (indicated in the table below). The injection mix was done with 4% phenol red, and 1nl was injected at a 1-cell stage. The amount of Control MO injected was the same as the total amount of experimental MO injected.

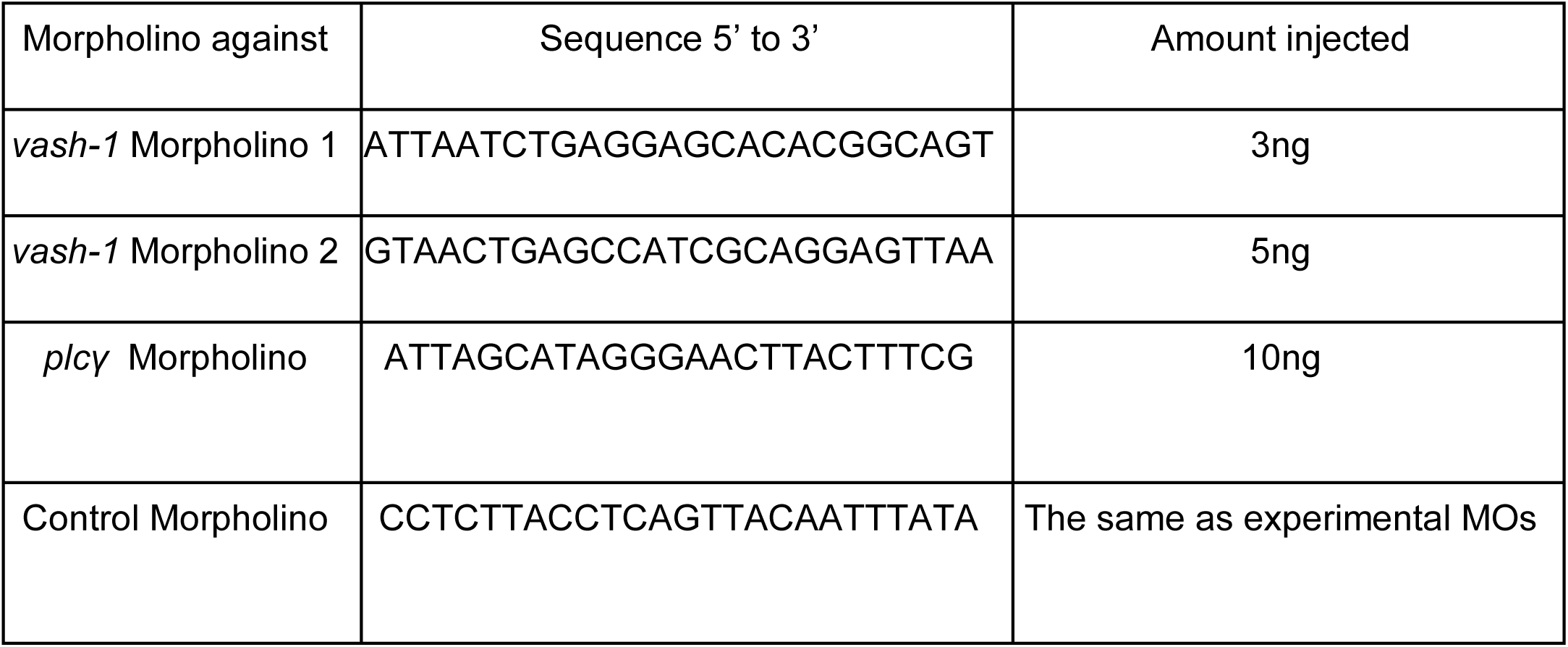

### Rescue experiment

*vash-1* full length cDNA (EMSEMBL ENSDART00000143819.3) was designed and synthesised by GenScript. TA overhangs were added by incubating the insert for 10 min at 72 °C with 50mM DNTPs and Taq polymerase (NEB), in order to perform TA cloning into a PCR4 TOPO vector (ThermoFisher). Integration was confirmed by sequencing. In order to transcribe the cDNA into RNA, the construct was linearised by NotI and the cDNA was transcribed for 4 hours by Megascript kit (LifeTechnologies) using the T3 promoter. The RNA was purified by Lithium-Chloride precipitation to a concentration of 500ng/μl. Aliquots were kept at −80°C until further usage. For rescue experiments, 1nl of 150pg RNA was injected, in embryos already injected with MOs against *vash-1* or control.

## RESULTS

### *Vasohibin-1* is highly expressed in zebrafish endothelial cells *in vivo*

To study the role of Vash-1 driven microtubule detyrosination in angiogenesis in zebrafish, we first investigated the conservation of zebrafish and human variants of Vash-1 (Fig. 1A,B) with CLUSTAL O (Uniprot). The analysis predicted 65,24% similarity between zebrafish and human Vash-1 (Fig. 1A), including the previously identified residues necessary for tubulin tyrosine and glutamate recognition, and the catalytic active site Cys-His-Ser triad (Adamopoulos et al., 2019; Nieuwenhuis et al., 2017; Sanchez-Pulido & Ponting, 2016) (Fig. 1B). RT-qPCR on RNA extracted from ECs sorted from *Tg[fli1a:nEGFP]*_*y7*_ embryos at 24 and 48 hours post fertilization (hpf) showed dynamic expression of *vash-1* during zebrafish development. During the sprouting phase (24hpf), *vash-1* expression was 5-7 times higher in endothelial than in non-ECs, decreasing at 48 hpf (Fig. 1C-D). Although these results are not significant, they were independently confirmed with a second primer set.

**Figure 1.**
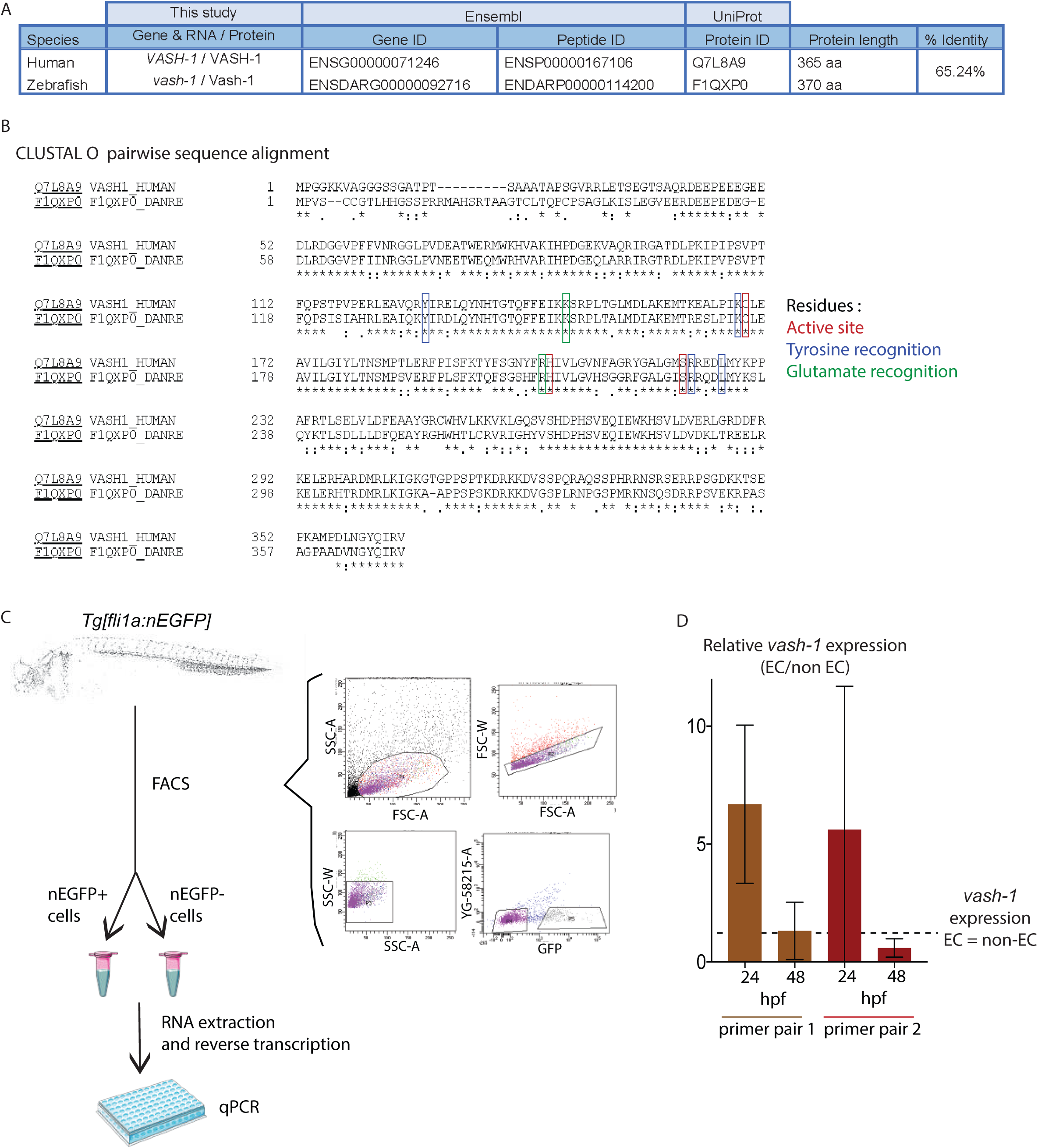
*vash-1* is highly expressed in zebrafish endothelium. **A-B** Aminoacid sequence alignment by CLUSTAL O (Uniprot) of human and zebrafish Vash-1 showing 65,1% sequence similarity (A), including in the residues necessary for tubulin detyrosination (B). **C-D** Analysis of *vash-1* expression in zebrafish embryos at 24 and 48hpf. FACS sorting and gating strategy for isolation of nEGFP+ and nEGFP-cells from *Tg[fli1a:nEGFP]*y7 embryos, their RNA extraction and reverse transcription leading to qPCR (C). *vash-1* expression, analysed by two independent primer pairs, show a tendency towards being enriched in the endothelium of zebrafish embryos at 24hpf (D). Data normalised to housekeeping genes. Each data point is an average of 3 technical replicates per sample, and each experiment has 3 biological replicates per developmental stage. Data is not significantly different, Mann-Whitney test.

### Vasohibin-1 carboxypeptidase function is conserved in zebrafish

A splice-site interfering morpholino (MO) targeting the intron3-exon4 boundary of nascent *vash-1* RNA (Fig. 2A) efficiently decreased the expression of *vash-1* in zebrafish embryos (Fig. 2B). Injections of MO against *vash-1* and control MO were performed in the *Tg[fli1ep:EGFP-DCX]* zebrafish line, selectively labelling endothelial microtubules by expression of GFP-tagged microtubule-binding protein doublecortin (Phng et al., 2015). Tubulin detyrosination was assessed by antibody staining against the glutamate residue of α-tubulin that becomes available after the tyrosine residue is removed by Vash-1 (Fig. 2C). Microtubules detected by this antibody are referred to as detyrosinated microtubules (dTyr). Whereas control embryos exhibited a high intensity array of detyrosinated tubulin in the neural tube, centrosomes, motoneurons (Fig. 2 D,E) and ECs (Fig. 2E’, arrowhead), *vash-1* knockdown (KD) embryos showed strongly diminished labelling of these structures (Fig. 2F,G). In particular, endothelial labelling was absent following *vash-1* KD (Fig. 2G’). Quantification of the ratio between dTyr tubulin and DCX (endothelial microtubules) signal intensities and of the detyrosination signal alone (Fig. 2H-I) confirmed that detyrosination of endothelial microtubules is significantly reduced in *vash-1* KD ISVs (dTyr/DCX= 0,24±0,01). in comparison with control ISVs (dTyr/DCX= 0,35±0,03) However, The DCX-GFP signal remained unchanged, suggesting that the overall microtubule cytoskeleton was not affected by *vash-1* KD (Fig. 2J).

**Figure 2.**
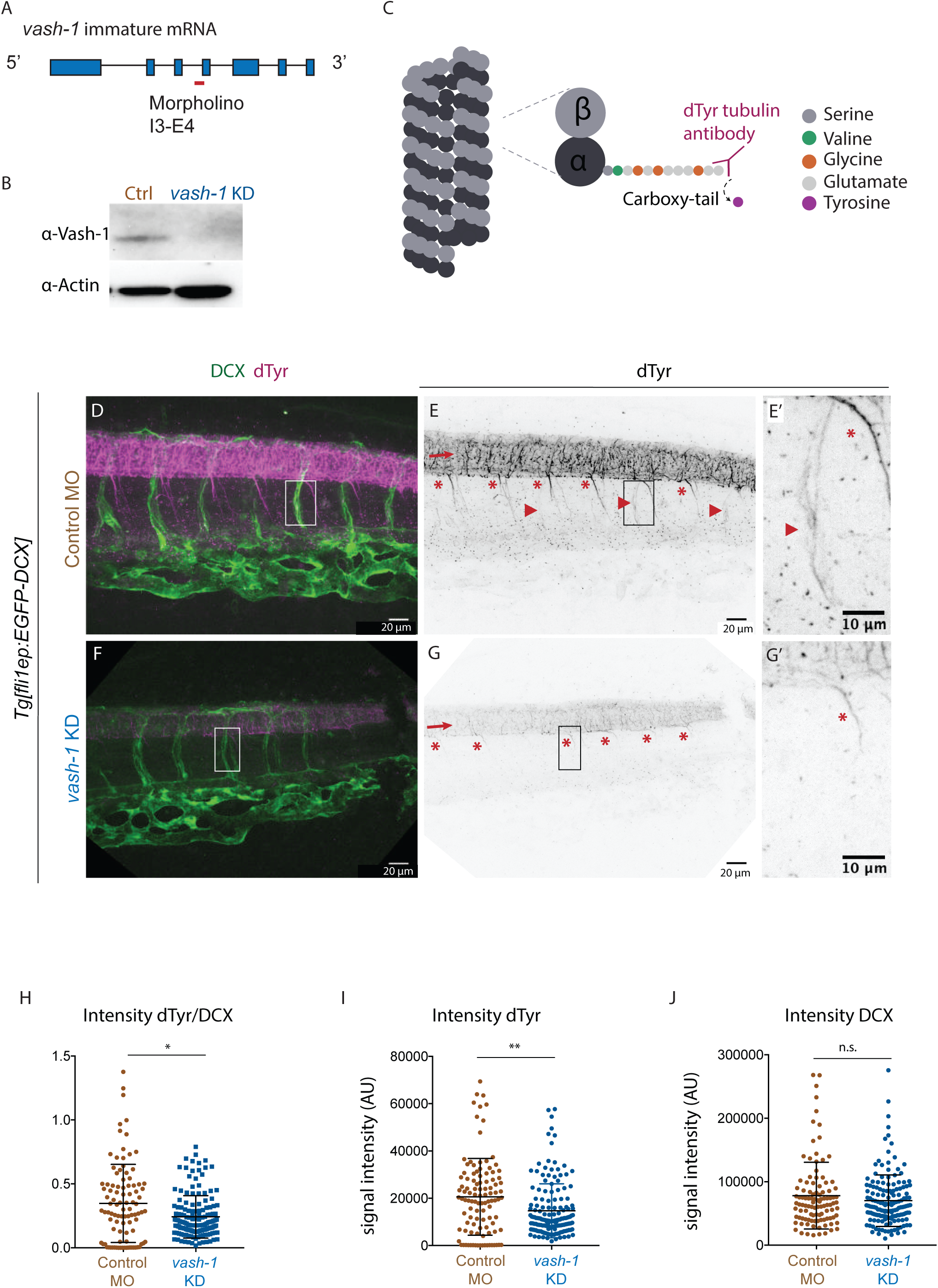
Endothelial microtubules are detyrosinated by Vash-1 in zebrafish. **A-B** Knockdown (KD) strategy using a morpholino (MO) targeting the intron3-exon4 (A) efficiently decreases Vash-1 protein levels as detected by Western blot (B). 48 hpf embryo lysate was used. **C** Mechanism of tyrosine cleavage from α-tubulin carboxy-terminus by Vash-1, resulting in detyrosinated tubulin (dTyr). Detyrosination exposes a glutamate residue accessible by a homemade antibody (Liao et al., 2019). **D-G** Immuno-stainings of 48 hpf *Tg[fli1ep:EGFP-DCX]* embryos detect detyrosinated microtubules (referred as dTyr, D,F,E,G) and GFP-labelled microtubules (referred as DCX, D,F). G,G’ shows quantification of dTyr signal upon *vash-1* KD compared with the control MO injected sibling embryos (E,E’). Arrows (E,G) indicate neural tube, with typically detyrosinated microtubules (E), reduced upon *vash-1* KD (G). Arrow heads indicate endothelial detyrosinated microtubules, only present in control embryos (E,E‘). Asterisks (E,G) indicate motoneurons exhibiting high dTyr signal in control embryos (E,E‘) and decreased dTyr signal in *vash-1* KD embryos (G,G’). **H-J** Quantifications of DCX and dTyr signal intensity of each ISV (data points) in control and *vash-1* KD groups. H. Quantification of ratios between dTyr and DCX intensity signals. Quantification of the intensity of the dTyr tubulin signal (I) and the EGFP-DCX signal (J) confirm that the microtubule detyrosination decreases in the ISVs whereas the eGFP-DCX intensity is not altered. AU stands for arbitrary units. Each data point is one ISV, N=100-110 ISVs from 9 embryos were quantified, in control and *vash-1* KD groups. p-values from Mann-Whitney test. * p<0.05, **, p<0.01. n.s.=not significant.

### Vasohibin-1 regulates secondary sprouting through control of endothelial cell divisions

Live-imaging of sprouting angiogenesis in *Tg[kdrl-l:ras-Cherry, fli1a:nEGFP]* embryos showed comparable primary sprouting and intersegmental vessel (ISV) network formation in control and *vash-1* KD embryos (Fig. 3A-F). Secondary sprouting however progressed abnormally in *vash-1* KD embryos with 1.5 times more secondary sprouts (Fig. 3F,G. Control embryos with 2,8±0,6 and *vash-1* embryos with 4,4±0,3). The *vash- 1* KD secondary sprouts exhibited an increased number of ECs (Fig. 3H,J) compared to control secondary sprouts (Fig. 3I). Quantifying nuclear divisions from movies acquired between 30 and 70 hpf at 15 minute time resolution revealed 3 times more nuclear divisions in the *vash-1* KD secondary sprouts than in controls (Fig. 3K). Furthermore, the transient 3-way connections (Fig. 3L-M) that regularly form during vein/lymphatic development (Geudens et al., 2019) took on average 1.5 times longer to resolve (Fig. 3N, control takes 6,0h±2,3 and vash-1 KD takes 11,3h±1,9). These results suggest a potential defect in the vasculature that forms by secondary sprouting.

**Figure 3.**
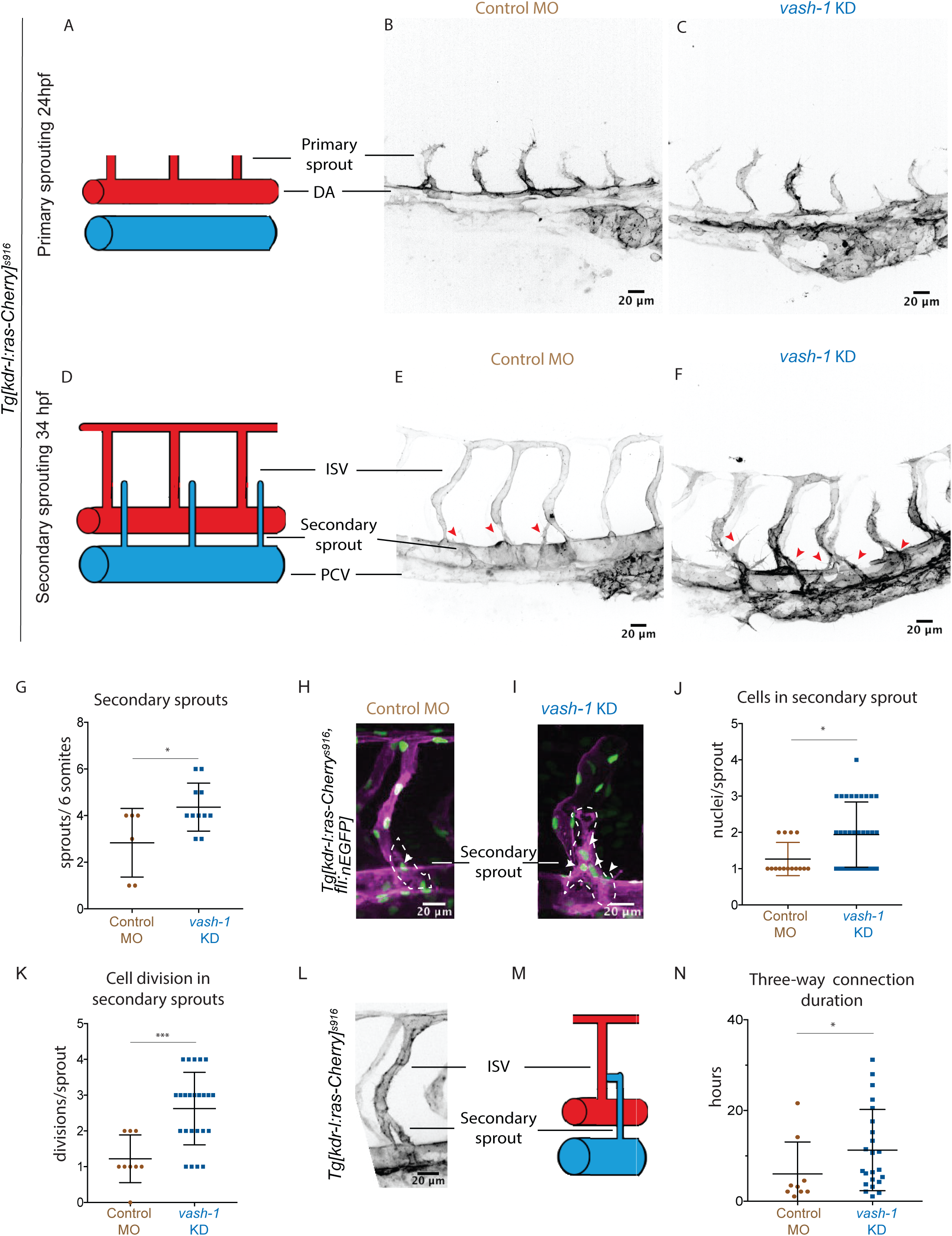
Vash-1 deficient embryos exhibit aberrant secondary sprouting and increased cell proliferation. **A-G** Primary and secondary sprouting in control and *vash-1* KD *Tg[kdr-l:ras-Cherry]s916* zebrafish embryos, with labelled blood vessels. Primary sprouting occurs from the dorsal aorta (DA) at 24–30 hours post fertilization (hpf) (A,B,C) and form inter-segmental vessels (ISVs) (D,E,F). Secondary sprouting occurs from the posterior cardinal vein (PCV) at 34-40 hpf (D,E,F). Red arrowheads label secondary sprouts (E,F). G. Quantifications of number of secondary sprouts per 6 somites. Data points come from 11 embryos for *vash-1* KD group, and for 6 control embryos. Mann-Whitney test, * = p<0.05. **H-K** Secondary sprouts in *Tg[kdr-l:ras-Cherrys916,fli1a:nEGFPy7]* embryos, with membrane- and nuclei-labelled endothelial cells (H,I) and quantification of endothelial nEGFP labelled nuclei in secondary sprouts (J) and number of nuclear divisions in the secondary sprout (K), from 30 to 70 hpf. White arrowheads indicate nuclei in secondary sprouts (marked by dashed line). N=9 ISVs from 4 control embryos, n=24 ISVs from 7 vash-1 KD embryos; P<0.0002 with t-test, ***= p<0.0002. **L-N** A transient 3-way connection is formed when the secondary sprout connects with the primary ISV and lumenises (L), schematically illustrated in M. N. Quantification of 3-way connection duration in control and *vash-1* MO injected embryos. Data points come from 28 ISVs from 9 embryos for *vash-1* KD and 11 ISVs from 5 embryos for control groups. p-value was calculated using Mann-Whitney test, * = p<0.05.

### Endothelial microtubules are selectively detyrosinated in secondary sprouts

Immunostaining against detyrosinated tubulin (dTyr) in fixed, uninjected *Tg[fli1ep:EGFP-DCX]* embryos at 24 and 34 hpf (Fig. 4A-F) revealed that, although the microtubules of the primary sprouts showed little to no staining (Fig. 4B,C,C’), the microtubules of the vast majority of secondary sprout cells were strongly labelled (Fig. 4E,F,F’). Quantification of fluorescence signal in endothelial microtubules from secondary sprouts was challenging due to the overlap of ECs from the primary sprouts. To mitigate this, we took advantage of the fact that zebrafish *phospholipase C-γ* (*plcγ)* morphants, that exhibit no primary sprouting and subsequent arterial ISV formation since the VEGFR2-Plcγ-signalling pathway is activated and necessary only for primary spouting (Hogan, Bos, et al., 2009). Secondary sprouting occurs normally in these embryos, and the absence of ISVs facilitates the measurement of signal intensities in secondary sprouts (Fig. 4G-I). Quantification of the signal intensity ratio between detyrosinated tubulin and DCX (endothelial microtubules) confirmed a 3-fold higher signal in secondary sprouts of *plcγ* morphants than in primary sprouts of control morphants (Fig. 4J. primary sprouts show dTyr signal intensity of 1,1±0,07 and secondary sprouts of 2,7±0,35).

**Figure 4.**
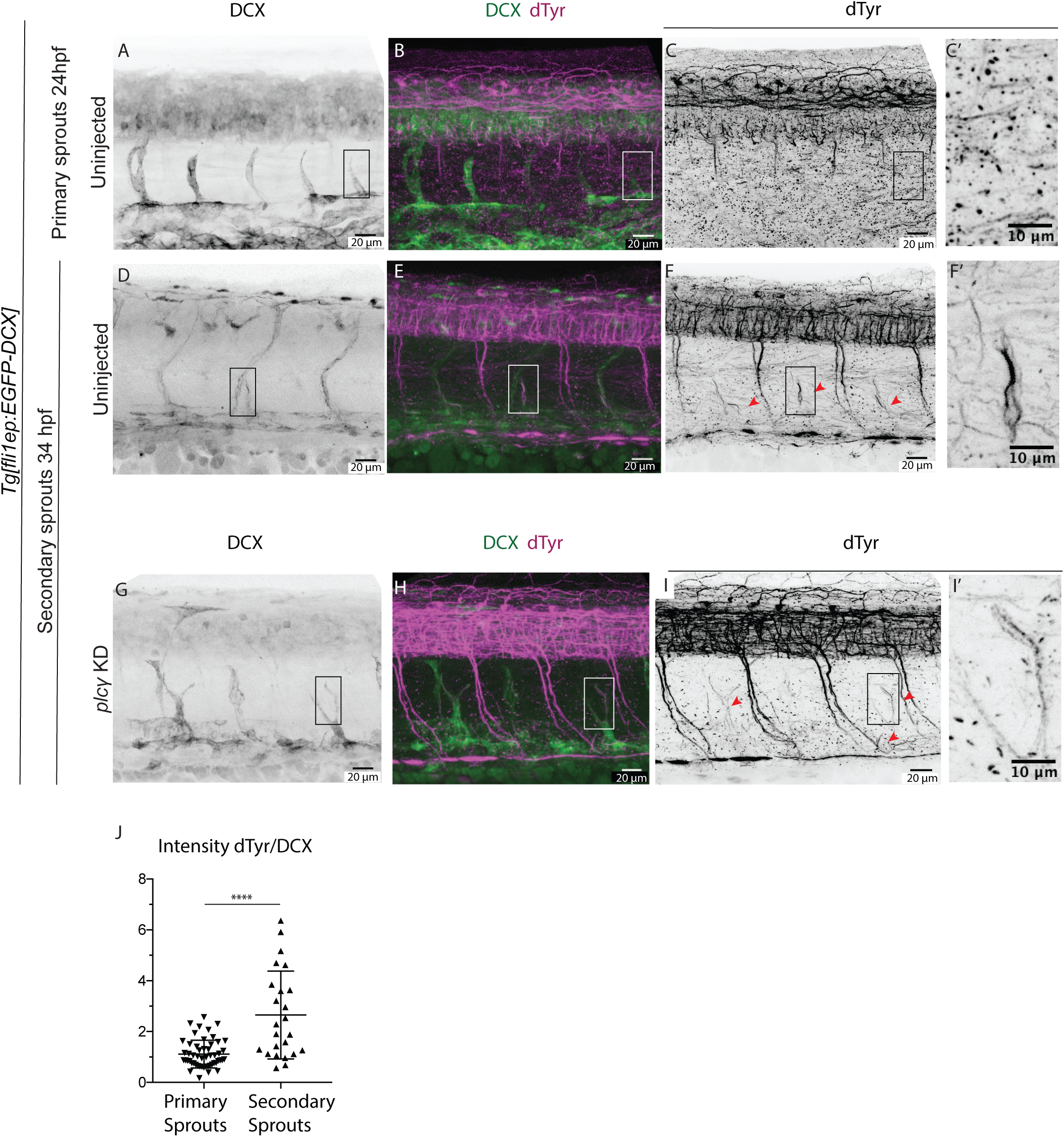
Microtubules of secondary sprouts are selectively highly detyrosinated. **A-I** Immunostainings using antibody detecting the glutamate aminoacid of detyrosinated microtubules (dTyr) during primary (A-C) and secondary (D-F) sprouting in uninjected *Tg[fli1ep:EGFP-DCX]* embryos, labelling all endothelial microtubules (DCX). C’, F’, I’, magnification from indicated region in C,F,I, respectively. *Plcγ* KD embryos (G-I) show reduced primary sprouting, facilitating the visualization and quantification of dTyr signal specifically in secondary sprouts. **J** Quantification of the proportion of dTyr/DCX signal intensity in primary sprouts of control morpholino (MO) injected embryos, and secondary sprouts from *Plcγ* MO injected embryos. N= 37 control primary sprouts, and N= 24 *Plcγ* KD secondary sprouts. p-value was calculated using Mann-Whitney test. *****<0.00001.

### Vasohibin-1 regulates formation of trunk lymphatic vasculature

Closer examination of *Tg[fli1a:EGFPy1,kdr-l:ras-Cherry]* embryos further revealed defects in the formation of lymphatic structures. At 52 hpf, the number of PLs emerging as lymphatic progenitors at the horizontal myoseptum was dramatically reduced in *vash-1* KD compared to control embryos (Fig. 5A-C). In addition, PLs in *vash-1* morphants frequently appeared connected to the venous ISVs, unlike in sibling controls (Fig. 5 D). Occasionally at 52 hpf and later in development at 4 dpf, we observed *kdrl-mCherry* positive, lumenised connections between ISVs, formed in the horizontal myoseptum (Fig. 5B,F,F’, arrow heads), suggesting they are aberrant vascular shunts. PLs are transient structures that normally form by sprouting of a subset of the secondary sprouts (Hogan, Bos, et al., 2009; Koltowska et al., 2015; Yaniv et al., 2006). From their initial position at the horizontal myoseptum, they migrate ventrally to shape the main trunk lymphatic vessel, the thoracic duct (TD) (Bussmann et al., 2010). The TD can be detected as an *eGFP* positive and *mCherry* negative vessel that locates between the DA and the PCV at 4 dpf. Control embryos showed almost complete TD formation along the trunk, whereas in *vash-1* morphants the TD was mostly absent (Fig. 5 E’,F’’). Quantification revealed a significant reduction in the percentage of somites with a TD fragment in *vash-1* KD embryos (20%±4,18) compared to controls (78%±4,4) (Fig. 5 G).

**Figure 5.**
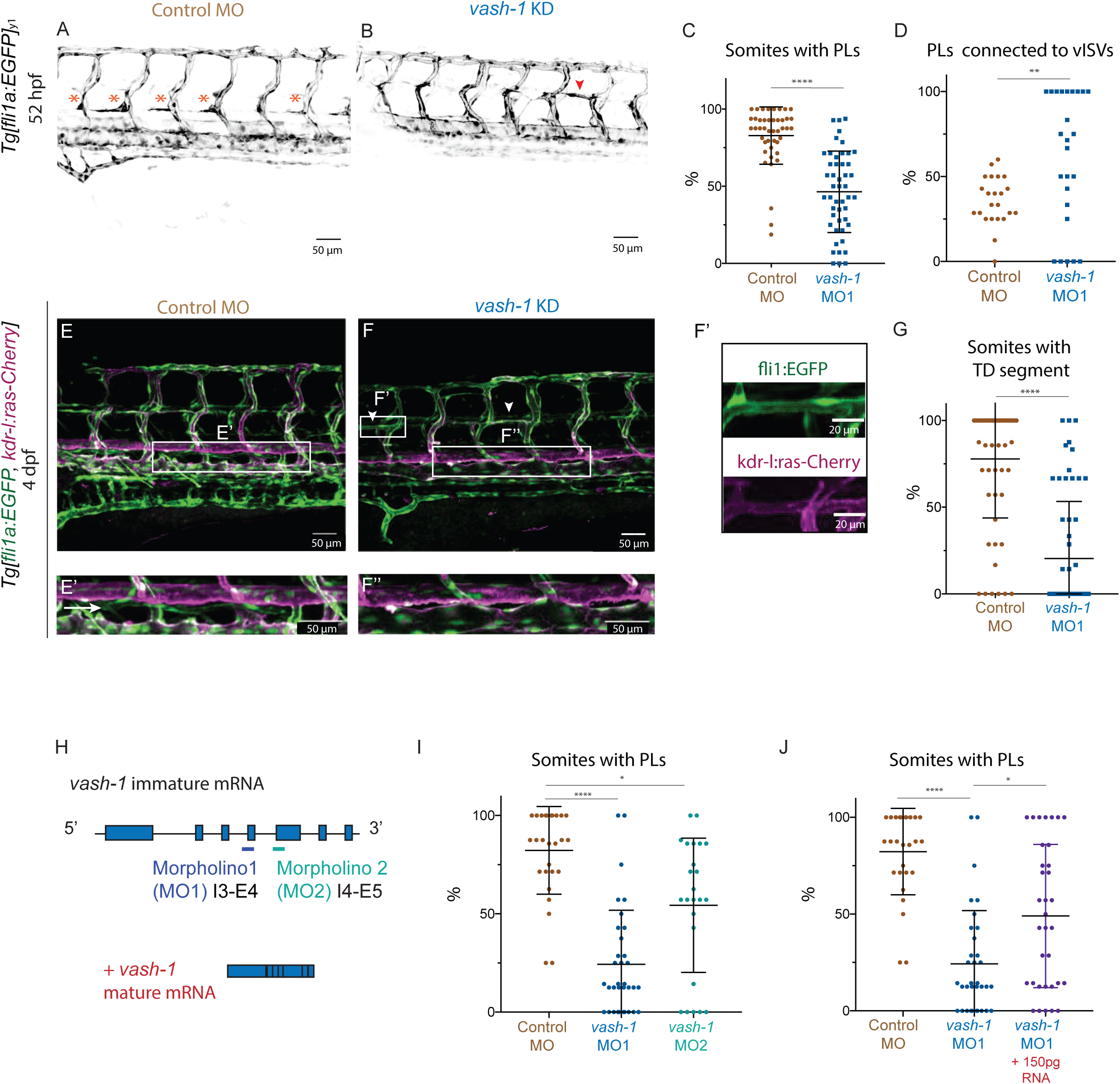
Vash-1 regulates formation of trunk lymphatic vasculature. **A-D** *Tg[fli1a:EGFP]y1* labels parachordal lymphangioblasts (PLs) of control (A) and *vash-1* KD embryos (B) at 52 hpf. Asterisks indicate PLs (A), arrowhead indicate putative ISV-to-ISV connection (B). C Quantification of the number of somites with PLs in embryos injected with control and *vash-1* MO. Quantification of percentage of PLs connected to a venous ISV (D). The amount of somites quantified varied from 6-8, n= 10-12 embryos per experimental group, minimum of 3 biological replicates per quantification, p-value was calculated using Mann-Whitney test. ****, p<0.0001, **=<0.02. **E-G** Zebrafish trunk of 4 dpf *Tg[kdr-l:ras-Cherrys916,fli1a:EGFPy1]* zebrafish embryos. Arrow heads indicate GFP and mCherry positive putative ISV-to-ISV connections in *vash-1* KD embryos (F,F’). The main axial lymphatic in the zebrafish trunk – thoracic duct (TD) - is GFP-positive, mCherry-negative (E,E’, labelled by an arrow), absent in the *vash-1* KD embryo (F,F‘’). **G** Quantification of the number of somites with TD. For all quantifications (C,D,G), the amount of somites quantified varied from 6-8, n= 10-12 embryos per experimental group, minimum of 3 biological replicates per quantification, p-value was calculated using Mann-Whitney test. ****, p<0.0001,**=<0.02. **H-J** Validation and quantification of KD experiment include confirmation of the phenotype with a second MO targeting a different splicing region of the *vash-1* immature mRNA and rescue of the phenotype with mature mRNA lacking the MO targeting region (H). Quantification of somites with PLs upon injection of a second MO (MO2) targeting *vash-1* (I) and co-injection of mature *vash-1* mRNA in *vash-1* MO1 injected embryos (J). For all groups, 22-30 embryos, and 7-8 somites quantified. p-value was calculated using Kruskal-Wallis test. ****, p<0.0001, **=<0.02, * = p<0.05.

The observation of the reduced number of PLs in the *vash-1* morphants was confirmed by a second MO targeting a different splicing region of nascent *vash-1* mRNA (Fig. 5H,I). Finally, we were able to partially rescue the loss of PLs in *vash-1* morphants by co-injection with *VASH-1* mRNA (Fig. 5J), supporting our hypothesis that the observed lymphatic defects are indeed caused by loss of Vash-1 function.

## DISCUSSION

Tubulin detyrosination is important for several differentiation events and developmental processes, such as myogenesis (Chang et al., 2002; Gundersen et al., 1989; Kerr et al., 2015) and neurogenesis (Aillaud et al., 2017; Erck et al., 2005), although the cellular mechanisms behind this are poorly understood. Vash-1, an angiogenesis inhibitor, was recently identified as the catalyst of this reaction (Aillaud et al., 2017; Nieuwenhuis et al., 2017). The cellular mechanisms behind the vascular phenotypes upon Vash-1 loss- and gain-of-function are not well understood.

In this study, we identified Vash-1-mediated microtubule detyrosination as a cellular mechanism of relevance for the control of EC sprouting from the PCV and the subsequent formation of lymphatic vessels in the zebrafish trunk. We showed that microtubules in secondary sprouts are particular in that they are selectively highly detyrosinated. We further show that *vash-1* depletion reduces the level of microtubule detyrosination and negatively affects secondary sprouting. Upon *vash-1* KD, supernumerous secondary sprouts emerge from the PCV. These sprouts contain more endothelial nuclei, as ECs divide almost twice as often as in control conditions. These results suggest that Vash-1-driven microtubule detyrosination limits excessive venous EC sprouting and proliferation during lympho-venous development in zebrafish.

Although the cellular mechanisms of primary and secondary sprouting in zebrafish appear very similar, with tip cell selection and guided migration and stalk cell proliferation, secondary sprouting utilises alternative signalling pathways and entails a unique specification step that establishes both venous ISVs and lymphatic structures. Secondary sprouting in zebrafish is triggered by Vegfc and activation of its receptor Flt4 in a subpopulation of ECs in the PCV (Bussmann et al., 2010; Hogan, Herpers, et al., 2009) that increase *prox1* expression prior to a cell division (Koltowska et al., 2015; Nicenboim et al., 2015). This cell division is thought to lead to asymmetric identity acquisition by the daughter cells, as one cell remains in the PCV while the other cell sprouts from the PCV to form a venous ISV (vISV) or become a lymphatic progenitor (Koltowska et al., 2015; Nicenboim et al., 2015). The specification of vISVor lymphatic progenitor is influenced by the levels of *prox1* expression, with high expression thought to favour lymphatic specification by reinforcing *flt4* expression (Koltowska et al., 2015). The observed increase of secondary sprouts in *vash-1* morphants, as well as the supernumerous EC numbers within these sprouts and the reduced numbers of somites with PLs, are consistent with defects in both the specification of daughter cells in the PCV and the specification of secondary sprouts into venous or lymphatic fates.

Intriguingly, microtubule detyrosination has been shown to influence mitotic and meiotic spindle (Akera et al., 2017; Barisic et al., 2015; Liao et al., 2019), with emerging evidence for a potential role in asymmetric cell fate (Akera et al., 2017; Laloraya, 2018). Given the numerous examples of acquisition of different fates and/or behaviours by daughter cells after cell division in various developing tissues (Boone & Doe, 2008; G. et al., 2016; Yu et al., 2006), including ECs (G. et al., 2016; Koltowska et al., 2015; Nicenboim et al., 2015), it is tempting to speculate that the lack of detyrosinated microtubules interferes with this process during initiation and progression of secondary sprouting, such that endothelial daughter cells retain their undifferentiated state and fail to specialise into lymphatic ECs (LECs). Failure to differentiate under activated conditions is frequently associated with continued proliferation, also in ECs (Benedito et al., 2008; Siekmann & Lawson, 2007). A recent study reported a similar increase in secondary sprout numbers in the *nova2* zebrafish mutants (Baek et al., 2019). Nova2 acts as a splicing factor that limits Vegfc/Flt4 signalling. Its absence causes premature and excessive formation of sprouting venous EC that fail to fully differentiate into functional lymphatic structures due to an increase of Vegfc/Flt4/Mapk signalling and consequently persistent proliferation.

According to current understanding, the decisive fate determining step in the formation of the lymphatic system in the zebrafish trunk is the acquisition of a *prox1a* high expression status in the cells of the PCV and reinforcement of the *flt4* signalling in a majority of lymphatic precursor cells. These cells emerge from the PCV as secondary sprouts to form first the PLs and subsequently expand to form the TD and other trunk lymphatic vessels (Koltowska et al., 2015; Nicenboim et al., 2015). Recent work demonstrated that a secondary sprout either contribute to remodelling a pre-existing ISV into a vein, or form a PLs (Geudens et al., 2019). In addition, a single secondary sprout is able to simultaneously form a vISV and give rise to lymphatic progenitors, which can be seen emerging from a remodelled vISVs (Geudens et al., 2019; Isogai et al., 2003). Unlike in control siblings, the few PLs forming in *vash-1* morphants are found associated with venous ISVs and often have a more vascular appearance, suggesting a defect in lymphatic specification. Additionally, the number of PLs present appears insufficient to form the trunk lymphatic vasculature. Together these findings suggest that Vash-1 is of particular importance to control lymphangiogenesis.

Vash-1 is highly conserved across eukaryotes, especially the three amino-acids that constitute a non-canonical Cys-His-Ser catalytic triad active site (Sanchez-Pulido & Ponting, 2016) that are necessary for the tubulin detyrosination function (Adamopoulos et al., 2019; Nieuwenhuis et al., 2017). Human and murine VASH-1 protein variants induce similar effects in angiogenesis assays when applied ectopically (Watanabe et al., 2004), suggesting similar functions across these species. Our pairwise sequence analysis indicates that the residues necessary for tubulin recognition and detyrosination (Adamopoulos et al., 2019; Nieuwenhuis et al., 2017) are conserved between human and zebrafish versions of Vash-1, suggesting conserved carboxypeptidase function. The fact that we could observe absence of microtubule detyrosination upon *vash-1* KD, and rescue the phenotype with a human variant of *VASH-1* mRNA further supports this hypothesis. Whilst earlier studies using gain-of-function approaches reported anti-lymphangiogenic activities of ectopically added VASH-1 in the mouse cornea and in tumour conditions (Heishi et al., 2010), our loss-of-function analysis in zebrafish suggests a selective pro-lymphangiogenic function of Vash-1 by supporting lympho-venous specification. Although this appears contradictory at first sight, a possible influence on Vegfc-mediated effects may explain the discrepancy. Whereas an acute inhibition of Vegfc signalling in existing lymphatic structures such as cornea and tumour inhibits further lymphatic growth, excessive Vegfc signalling during initial sprouting from the PCV interferes with lymphatic differentiation. This occurs in the *nova2* mutant (Baek et al., 2019), and potentially also in the absence of Vash-1. Future work will need to establish whether and how Vash-1 mediated detyrosination of endothelial microtubules restricts excessive Vegfc signalling, in some or all daughter cells during formation of secondary sprouts from the PCV.

In conclusion, our work identifies Vash-1 as a conserved tubulin carboxypeptidase in zebrafish, with a selective role in secondary sprouting and LEC formation. We propose that a balance of the number of ECs sprouting from the cardinal vein is maintained by Vash-1 via microtubule detyrosination and consequent EC divisions and that Vash-1 has a role in LEC specification and formation in the zebrafish trunk.

## ACKNOWLEDGEMENTS

We thank members of the Gerhardt lab for the discussions, in particular to Dr. André Rosa for the *Tg[fli1a:nEGFP]*_*y7*_ embryo picture in figure 1. We thank Simone Reber and Carsten Janke for important conceptual discussions. We gratefully acknowledge excellent support by Robby Fechner and the staff of the MDC aquatic facility.

## REFERENCES

Abe, M., & Sato, Y. (2001). cDNA microarray analysis of the gene expression profile of VEGF-activated human umbilical vein endothelial cells. Angiogenesis, 4(4), 289–298.

Adamopoulos, A., Landskron, L., Heidebrecht, T., Tsakou, F., Bleijerveld, O. B., Altelaar, M., Nieuwenhuis, J., Celie, P. H. N., Brummelkamp, T. R., & Perrakis, A. (2019). Crystal structure of the tubulin tyrosine carboxypeptidase complex VASH1–SVBP. Nature Structural & Molecular Biology, 26(7), 567–570.

Aillaud, C., Bosc, C., Peris, L., Bosson, A., Heemeryck, P., Dijk, J. Van, Friec, J. Le, Boulan, B., Vossier, F., Sanman, L. E., Syed, S., Amara, N., Couté, Y., Lafanechère, L., Denarier, E., Delphin, C., Pelletier, L., Humbert, S., Bogyo, M., & Andrieux, A. (2017). Vasohibins/SVBP are tubulin carboxypeptidases (TCPs) that regulate neuron differentiation. 1453(December), 1448–1453.

Akera, T., Chmátal, L., Trimm, E., Yang, K., Aonbangkhen, C., Chenoweth, D. M., Janke, C., Schultz, R. M., & Lampson, M. A. (2017). Spindle asymmetry drives non-Mendelian chromosome segregation. Science, 358(November), 668–672.

Aleström, P., D’Angelo, L., Midtlyng, P. J., Schorderet, D. F., Schulte-Merker, S., Sohm, F., & Warner, S. (2019). Zebrafish: Housing and husbandry recommendations. Laboratory Animals, 0(0), 1–12.

Baek, S., Oh, T. G., Secker, G., Sutton, D. L., Okuda, K. S., Paterson, S., Bower, N. I., Toubia, J., Koltowska, K., Capon, S. J., Baillie, G. J., Simons, C., Muscat, G. E. O., Lagendijk, A. K., Smith, K. A., Harvey, N. L., & Hogan, B. M. (2019). The Alternative Splicing Regulator Nova2 Constrains Vascular Erk Signaling to Limit Specification of the Lymphatic Lineage. Developmental Cell, 49(2), 279–292.e5.

Barisic, M., Silva E Sousa, R., Tripathy, S. K., Magiera, M. M., Zaytsev, A. V., Pereira, A. L., Janke, C., Grishchuk, E. L., & Maiato, H. (2015). Microtubule detyrosination guides chromosomes during mitosis. Science, 348(6236), 799–803.

Belmont, L. D., Hyman, A. A., Sawin, K. E., & Mitchison, T. J. (1990). Real-time visualization of cell cycle-dependent changes in microtubule dynamics in cytoplasmic extracts. Cell, 62(3), 579–589.

Benedito, R., Trindade, A., Hirashima, M., Henrique, D., Da Costa, L. L., Rossant, J., Gill, P. S., & Duarte, A. (2008). Loss of Notch signalling induced by Dll4 causes arterial calibre reduction by increasing endothelial cell response to angiogenic stimuli. BMC Developmental Biology, 8, 1–15.

Boone, J. Q., & Doe, C. Q. (2008). Identification of Drosophila type II neuroblast lineages containing transit amplifying ganglion mother cells. Developmental Neurobiology, 68(9), 1185–1195.

Bussmann, J., Bos, F. L., Urasaki, A., Kawakami, K., Duckers, H. J., & Schulte-Merker, S. (2010). Arteries provide essential guidance cues for lymphatic endothelial cells in the zebrafish trunk. Development, 137(16), 2653–2657.

Chang, W., Webster, D. R., Salam, A. A., Gruber, D., Prasad, A., Eiserich, J. P., & Chloë Bulinski, J. (2002). Alteration of the C-terminal amino acid of tubulin specifically inhibits myogenic differentiation. Journal of Biological Chemistry, 277(34), 30690–30698.

Coxam, B., Sabine, A., Bower, N. I., Smith, K. A., Pichol-thievend, C., Skoczylas, R., Astin, J. W., Frampton, E., Jaquet, M., Philip, S., Parton, R. G., Harvey, N. L., Petrova, T.V, Schulte-, S., Francois, M., & Hogan, B. M. (2014). Pkd1 regulates lymphatic vascular morphogenesis during development. Cell Rep., 7(3), 623–633.

Dunn, S., Morisson, E. E., Liverpool, T. B., Molina-París, C., Cross, R. A., Alonso, M. C., & Peckham, M. (2008). Differential trafficking of Kif5c on tyrosinated and detyrosinated microtubules in live cells. Journal of Cell Science, 121(7), 1085–1095.

Erck, C., Peris, L., Andrieux, A., Meissirel, C., Gruber, A. D., Vernet, M., Schweitzer, A., Saoudi, Y., Pointu, H., Bosc, C., Salin, P. A., Job, D., & Wehland, J. (2005). A vital role of tubulin-tyrosine-ligase for neuronal organization. Proceedings of the National Academy of Sciences of the United States of America, 102(22), 7853–7858.

G., C., Harrington, K., Lovegrove, H., Page, D., Chakravartula, S., Bentley, K., & Herbert, S. (2016). Asymmetric division coordinates collective cell migration in angiogenesis. Nature Cell Biology, 18(12), accepted.

Gebala, V., Collins, R., Geudens, I., Phng, L.-K., & Gerhardt, H. (2016). Blood flow drives lumen formation by inverse membrane blebbing during angiogenesis in vivo. Nature Cell Biology, advance on(February).

Geudens, I., Coxam, B., Alt, S., Gebala, V., Vion, A.-C., Meier, K., Rosa, A., & Gerhardt, H. (2019). Artery-vein specification in the zebrafish trunk is pre-patterned by heterogeneous Notch activity and balanced by flow-mediated fine-tuning. Development, 146(16), dev181024.

Geudens, I., Herpers, R., Hermans, K., Segura, I., Ruiz De Almodovar, C., Bussmann, J., De Smet, F., Vandevelde, W., Hogan, B. M., Siekmann, A., Claes, F., Moore, J. C., Silvia Pistocchi, A., Loges, S., Mazzone, M., Mariggi, G., Bruyère, F., Cotelli, F., Kerjaschki, D., … Dewerchin, M. (2010). Role of delta-like-4/notch in the formation and wiring of the lymphatic network in zebrafish. Arteriosclerosis, Thrombosis, and Vascular Biology, 30(9), 1695–1702.

Gore, A. V, Monzo, K., Cha, Y. R., Pan, W., & Weinstein, B. M. (2016). Vascular Development in the Zebrafish. 1–22.

Gundersen, G. G., Khawaja, S., & Bulinski, J. C. (1989). Generation of a stable, posttranslationally modified microtubule array is an early event in myogenic differentiation. Journal of Cell Biology, 109(5), 2275–2288.

Hayot, C., Farinelle, S., De Decker, R., Decaestecker, C., Darro, F., Kiss, R., & Van Damme, M. (2002). In vitro pharmacological characterizations of the anti-angiogenic and anti-tumor cell migration properties mediated by microtubule-affecting drugs, with special emphasis on the organization of the actin cytoskeleton.

Heishi, T., Hosaka, T., Suzuki, Y., Miyashita, H., Oike, Y., Takahashi, T., Nakamura, T., Arioka, S., Mitsuda, Y., Takakura, T., Hojo, K., Matsumoto, M., Yamauchi, C., Ohta, H., Sonoda, H., & Sato, Y. (2010). Endogenous angiogenesis inhibitor vasohibin1 exhibits broad-spectrum antilymphangiogenic activity and suppresses lymph node metastasis. American Journal of Pathology, 176(4), 1950–1958.

Hogan, B. M., Bos, F. L., Bussmann, J., Witte, M., Chi, N. C., Duckers, H. J., & Schulte-Merker, S. (2009). Ccbe1 is required for embryonic lymphangiogenesis and venous sprouting. Nature Genetics, 41(4), 396–398.

Hogan, B. M., Herpers, R., Witte, M., Heloterä, H., Alitalo, K., Duckers, H. J., & Schulte-Merker, S. (2009). Vegfc/Flt4 signalling is suppressed by Dll4 in developing zebrafish intersegmental arteries. Development, 136(23), 4001–4009.

Hogan, B. M., & Schulte-Merker, S. (2017). How to Plumb a Pisces: Understanding Vascular Development and Disease Using Zebrafish Embryos. Developmental Cell, 42(6), 567–583.

Hosaka, T., Kimura, H., Heishi, T., Suzuki, Y., Miyashita, H., Ohta, H., Sonoda, H., Moriya, T., Suzuki, S., Kondo, T., & Sato, Y. (2009). Vasohibin-1 expression in endothelium of tumor blood vessels regulates angiogenesis. American Journal of Pathology, 175(1), 430–439.

Isogai, S., Lawson, N. D., Torrealday, S., Horiguchi, M., & Weinstein, B. M. (2003). Angiogenic network formation in the developing vertebrate trunk. Development, 130(21), 5281–5290.

Kern, J., Steurer, M., Gastl, G., Gunsilius, E., & Untergasser, G. (2009). Vasohibin inhibits angiogenic sprouting in vitro and supports vascular maturation processes in vivo. BMC Cancer, 9, 284.

Kerr, J. P., Robison, P., Shi, G., Bogush, A. I., Kempema, A. M., Hexum, J. K., Becerra, N., Harki, D. A., Martin, S. S., Raiteri, R., Prosser, B. L., & Ward, C. W. (2015). Detyrosinated microtubules modulate mechanotransduction in heart and skeletal muscle. Nature Communications, 6.

Kimmel, C. B., Ballard, W. W., Kimmel, S. R., Ullmann, B., & Schilling, T. F. (1995). Stages of embryonic development of the zebrafish. Developmental Dynamics : An Official Publication of the American Association of Anatomists, 203(3), 253–310.

Kimura, H., Miyashita, H., Suzuki, Y., Kobayashi, M., Watanabe, K., Sonoda, H., Ohta, H., Fujiwara, T., Shimosegawa, T., & Sato, Y. (2009). Distinctive localization and opposed roles of vasohibin-1 and vasohibin-2 in the regulation of angiogenesis. Blood, 113(19), 4810–4818.

Kohli, V., Schumacher, J. A., Desai, S. P., Rehn, K., & Sumanas, S. (2013). Arterial and Venous Progenitors of the Major Axial Vessels Originate at Distinct Locations. Developmental Cell, 25(2), 196–206.

Koltowska, K., Lagendijk, A. K., Pichol-Thievend, C., Fischer, J. C., Francois, M., Ober, E. A., Yap, A. S., & Hogan, B. M. (2015). Vegfc Regulates Bipotential Precursor Division and Prox1 Expression to Promote Lymphatic Identity in Zebrafish. Cell Reports, 13(9), 1828–1841.

Kruczynski, A., Poli, M., Dossi, R., Chazottes, E., Berrichon, G., Ricome, C., Giavazzi, R., Hill, B. T., & Taraboletti, G. (2006). Anti-angiogenic, vascular-disrupting and anti-metastatic activities of vinflunine, the latest vinca alkaloid in clinical development. European Journal of Cancer, 42(16), 2821–2832.

Küchler, A. M., Gjini, E., Peterson-Maduro, J., Cancilla, B., Wolburg, H., & Schulte-Merker, S. (2006). Development of the Zebrafish Lymphatic System Requires Vegfc Signaling. Current Biology, 16(12), 1244–1248.

Laloraya, S. (2018). Asymmetric Tyrosination of Spindle Microtubules Facilitates Selfish Inheritance. Trends in Cell Biology, 28(6), 417–419.

Lamalice, L., Le Boeuf, F., & Huot, J. (2007). Endothelial cell migration during angiogenesis. Circulation Research, 100(6), 782–794.

Lawson, N. D., & Weinstein, B. M. (2002). In Vivo Imaging of Embryonic Vascular Development Using Transgenic Zebrafish. Developmental Biology, 248(2), 307–318.

Liao, S., Rajendraprasad, G., Wang, N., Eibes, S., Gao, J., Yu, H., Wu, G., Tu, X., Huang, H., Barisic, M., & Xu, C. (2019). Molecular basis of vasohibins-mediated detyrosination and its impact on spindle function and mitosis. Cell Research, June.

Nicenboim, J., Malkinson, G., Lupo, T., Asaf, L., Sela, Y., Mayseless, O., Gibbs-Bar, L., Senderovich, N., Hashimshony, T., Shin, M., Jerafi-Vider, A., Avraham-Davidi, I., Krupalnik, V., Hofi, R., Almog, G., Astin, J. W., Golani, O., Ben-Dor, S., Crosier, P. S., … Yaniv, K. (2015). Lymphatic vessels arise from specialized angioblasts within a venous niche. Nature, 522(7554), 56–61.

Nieuwenhuis, J., Adamopoulos, A., Bleijerveld, O. B., Mazouzi, A., Stickel, E., Celie, P., Altelaar, M., Knipscheer, P., Perrakis, A., Blomen, V. A., & Brummelkamp, T. R. (2017). Vasohibins encode tubulin detyrosinating activity. Science, 358(6369), 1453–1456.

Pfaffl, M. (2001). A new mathematical model for relative quantification in real-time RT-PCR. Nucleic Acids Research, 29(9), 2002–2007.

Phng, L.-K., Gebala, V., Bentley, K., Philippides, A., Wacker, A., Mathivet, T., Sauteur, L., Stanchi, F., Belting, H.-G., Affolter, M., & Gerhardt, H. (2015). Formin-Mediated Actin Polymerization at Endothelial Junctions Is Required for Vessel Lumen Formation and Stabilization. Developmental Cell, 32, 123–132.

Potente, M., Gerhardt, H., & Carmeliet, P. (2011). Basic and therapeutic aspects of angiogenesis. Cell, 146(6), 873–887.

Roman, B. L., Pham, V. N., Lawson, N. D., Kulik, M., Childs, S., Lekven, A. C., Garrity, D. M., Moon, R. T., Fishman, M. C., Lechleider, R. J., & Wienstein, B. M. (2002). Disruption of acvrl1 increases endothelial cell number in zebrafish cranial vessels. Development, 129(12), 3009–3019.

Ruijter, J. M., Ramakers, C., Hoogaars, W. M. H., Karlen, Y., Bakker, O., van den hoff, M. J. B., & Moorman, A. F. M. (2009). Amplification efficiency: Linking baseline and bias in the analysis of quantitative PCR data. Nucleic Acids Research, 37(6).

Sanchez-Pulido, L., & Ponting, C. P. (2016). Vasohibins: New transglutaminase-like cysteine proteases possessing a non-canonical Cys-His-Ser catalytic triad. Bioinformatics, 32(10), 1441–1445.

Schwartz, E. L. (2009). Antivascular actions of microtubule-binding drugs. Clinical Cancer Research, 15(8), 2594–2601.

Shi, Y.-W., Yuan, W., Wang, X., Gong, J., Zhu, S.-X., Chai, L.-L., Qi, J.-L., Qin, Y.-Y., Gao, Y., Zhou, Y.-L., Fan, X.-L., Ji, C.-Y., Wu, J.-Y., Wang, Z.-W., & Liu, D. (2016). Combretastatin A-4 efficiently inhibits angiogenesis and induces neuronal apoptosis in zebrafish. Scientific Reports, 6, 30189.

Shibuya, T., Watanabe, K., Yamashita, H., Shimizu, K., Miyashita, H., Abe, M., Moriya, T., Ohta, H., Sonoda, H., Shimosegawa, T., Tabayashi, K., & Sato, Y. (2006). Isolation and characterization of vasohibin-2 as a homologue of VEGF-inducible endothelium-derived angiogenesis inhibitor vasohibin. Arteriosclerosis, Thrombosis, and Vascular Biology, 26(5), 1051–1057.

Siegrist, S. E., & Doe, C. Q. (2007). Microtubule-induced cortical cell polarity. Genes and Development, 21(5), 483–496.

Siekmann, A. F., & Lawson, N. D. (2007). Notch signalling limits angiogenic cell behaviour in developing zebrafish arteries. Nature, 445(7129), 781–784.

Tozer, G. M., Kanthou, C., Parkins, C. S., & Hill, S. a. (2002). The biology of the combretastatins as tumour vascular targeting agents. International Journal of Experimental Pathology, 83(1), 21–38.

Vale, R. D. (2003). The molecular motor toolbox for intracellular transport. Cell, 112(4), 467–480.

Watanabe, K., Hasegawa, Y., Yamashita, H., Shimizu, K., Ding, Y., Abe, M., Ohta, H., Imagawa, K., Hojo, K., Maki, H., Sonoda, H., & Sato, Y. (2004). Vasohibin as an endothelium-derived negative feedback regulator of angiogenesis. Journal of Clinical Investigation, 114(7), 898–907.

Yaniv, K., Isogai, S., Castranova, D., Dye, L., Hitomi, J., & Weinstein, B. M. (2006). Live imaging of lymphatic development in the zebrafish. Nature Medicine, 12(6), 711–716.

Yu, F., Kuo, C. T., & Jan, Y. N. (2006). Drosophila Neuroblast Asymmetric Cell Division: Recent Advances and Implications for Stem Cell Biology. Neuron, 51(1), 13–20.

